# OSM-11 modulates salinity-stress tolerance in *Caenorhabditis elegans*

**DOI:** 10.1101/2025.11.20.689412

**Authors:** Pengchi Zhang, Beining Xue, Yusu Xie, Kunlin Li, Hanwen Yang, Peiqi Sun, Liusuo Zhang

**Author notes:** Co-first Authors (Pengchi Zhang, Beining Xue, Yusu Xie).

## Abstract

Most terrestrial animals exhibit narrow salinity tolerance compared to their marine counterparts. Previous studies identified *osm-11* (which encodes a Notch co-ligand) mutations as a driver of hyper-saline tolerance in *Caenorhabditis elegans*, but mechanistic insights remained unclear. This study employs RNA sequencing and CRISPR/Cas-9 genome editing to demonstrate that *osm-11* mutations enhance salinity stress resistance through up-regulation of fatty acid metabolism (*acdh-12*, *acs-17*) and cytochrome P450 pathways (*ugt-15*), while suppressing calcium signaling. Furthermore, we demonstrated that *acdh-12* mutation impairs salinity-stress tolerance by activating ferroptosis and mitophagy, accompanied by down-regulated oxidative phosphorylation and up-regulated autophagic pathways. Morphological observations show that mitochondrial fragmentation contributes to wild-type nematode mortality under high salinity, while enlarged lipid droplets in wild-types correlate with reduced β-oxidation gene expression (*dhs-28*, *daf-22*), whose knockout disrupts tolerance in mutants. These findings unravel the multi-pathway regulatory network of *osm-11*-mediated salinity tolerance, providing mechanistic insights for developing protective strategies against environmental salinity stressors impacting animal survival.

## Introduction

Recent sea level rise induced by global climate change could accelerate marine water intrusion into fertile soils - at the same time, excessive groundwater extraction and widespread use of synthetic fertilizers also increase soil salinity ^1^. Soil salinization has become an emerging threat to terrestrial ecosystems, impairing sustainable productions of agriculture and aquaculture ^2^. Studies have shown that TRPV channels and sodium channels played essential roles in osmoregulation in mammalian neurons, and the purinergic P2Y2 receptor was involved in renal osmoregulation ^3^. Osmoregulatory tissues and organs, such as gills in fish or kidneys in mammals, played important roles in response to hyposaline or hyper-saline environmental conditions. Additionally, the evolutionarily conserved cellular stress response (CSR) was utilized to cope with salinity stresses in fish ^4,5^.

Several osmoregulation mechanisms have been reported in invertebrates ^6^. Such mechanisms involve transcriptional and post-transcriptional regulation of genes in MAPK and insulin-like signaling pathways ^7–9^, ion transport-related genes encoding transient receptor potential cation channel and chloride channel ^10,11^, aquaporin water channel genes ^12^, and genes essential for the synthesis and transport of organic osmolytes- such as glycerol, myo-inositol, taurine, methylamines, as well as proline, sucrose, and trehalose ^6^.

*C. elegans* is a powerful biomedical invertebrate model used to investigate the mechanisms underlying animal salinity stress response and their adaptations ^11,13–15^. It was reported that in *C. elegans* ASH neurons triggered hyperosmotic avoidance behavior ^16,17^ while ALA neurons induced sleep-like quiescent under hyper-saline environment ^18^. Scientists have elucidated the pivotal roles of transmembrane proteins OCR-2, OSM-9, G protein ODR-3, and OSM-10 in the sensing and transduction of extracellular osmotic pressure in ASH neurons ^19^. In nematodes, the excretory system, epidermis, and intestine are all principal contributors to osmotic regulation ^20,21^.

Glycerol is the major organic osmolyte in *C. elegans* under hyperosmotic environments ^22^. In high-salinity environments, *C. elegans* rapidly induced the salinity-stress responses by upregulating *gpdh-1* (which encodes glycerol-3-phosphate dehydrogenase) to enhance glycerol synthesis ^10,11^. *gpdh-1* and *gpdh-2* are key glycerol synthesis enzymes in *C. elegans*, and *gpdh-1*;*gpdh-2* double mutants showed a significant decrease in glycerol accumulation, fertility and developmental delay in hyperosmotic environments ^22^. GPDH-2 was required for intergenerational osmotic protection and decreased maternal germline insulin-like signaling protected progeny from hyperosmotic stress ^23^. *flcn-1* mutants exhibited increased hyper-saline tolerance via AMPK mediated glycogen accumulation ^24^. Cytosolic sulfotransferase SSU-1 antagonized insulin-like signaling under hyperosmotic conditions ^25^. Mutations in the insulin receptor gene, *daf-2*, and its downstream *age-1* gene (which encodes PI3K) conferred hypertonic resistance in a DAF-16-dependent manner ^26,27^. Salinity-tolerance was severely impaired in *daf-2/age-1* mutants when both trehalose synthesis genes *tps-1/tps-2* were knocked down ^26–28^. In addition, knocking down either glycogen synthesis gene *gsy-1*, or glycogen phosphorylase gene *pygl-1*, significantly impaired salinity-tolerance in *daf-2* mutants^26^.

It was reported that mutations in *osm-7* and *osm-11* induced hyper-saline resistance in *C. elegans* ^29^, both genes function as co-ligands to activate Notch receptors LIN-12 and GLP-1 ^30^. *osm-7;osm-11* double mutants exhibited the same salinity-stress tolerance phenotype as *osm-7* ^29^, indicating that these genes regulate osmotic resistance through a shared pathway ^29^. OSM-11 was reported to regulate heat and oxidative stress responses in a *skn-1* dependent manner, while *osm-11* mutants promoted animal hyper-saline resistance independent of *skn-1*^31^. However, the molecular mechanism by which OSM-11 regulates osmotic resistance remains unclear.

In this study, we observed that *osm-11* mutants showed significantly increased hyper-saline tolerance compared to wild-type animals and dozens of other osmotic resistance mutants, suggesting a major role for *osm-11* in salinity-stress regulation. Using CRISPR/Cas9 genome editing, RNAi, comparative transcriptomics, and imaging, we revealed the mechanisms by which Notch co-ligand OSM-11 mediates salinity-stress tolerance in *C. elegans*. We generated several *osm-11* mutants with different alleles, confirming that mutations in *osm-11* conferred to tolerance to hyper-saline stress. Next, comparative transcriptomic analysis between *osm-11* mutants and wild-type N2 animals under different salinity conditions revealed a significantl up-regulattion of gene expression in KEGG pathways, such as fatty acid metabolism and cytochrome P450 in *osm-11* mutants, while calcium pathway genes (e.g., *ncx-1*, *ncx-2*, *ckr-2*, *cmk-1*, *gar-3*) were significantly down-regulated. Intriguingly, CRISPR/Cas9-mediated knock-out of candidate genes identified that mutations in *acdh-12, acs-17* and *ugt-15* significantly reduced salinity-stress tolerance in *osm-11* mutants. Furthermore, transcriptomic comparison between *osm-11* single mutants and *osm-11;acdh-12* double mutants showed significantly enhanced expression of mitochondria-related KEGG pathways (e.g., oxidative phosphorylation, citrate cycle/TCA cycle) in *osm-11* mutants. Salinity-stress tolerance in *osm-11* mutants also correlated with suppressed mitochondrial fragmentation. Additionally, RNAi knockdown experiments revealed that *gpx-1/gpx-7* depletion significantly reduced *osm-11* mutant survival under hyper-saline environment, while *dct-1* knockdown significantly improved survival of *osm-11;acdh-12*, indicating that *acdh-12* mutation impairs salinity tolerance in *osm-11* mutants via enhanced ferroptosis and mitophagy.

## Material and methods

### *C. elegans* strains

Worms were maintained on Nematode Growth Medium (NGM) plates seeded with *E. coli* OP50 at 20°C. Bristol N2 was utilized as the wild-type control. For strain information see Table S1.

### Sea salt NGM plates preparation

Sea salt NGM with concentrations of 3‰, 6‰, 9‰, 12‰, 15‰, and 30‰ were prepared by adding artificial sea salt (Instant Ocean Sea Salt) to 300ml of NGM, respectively. For details. see Section S1.

### Hyper-saline resistance assay

To assess the hyper-saline tolerance of mutants, synchronized L1 larvae were transferred into 35mm 30‰ salinity NGM treatment plates seeded with 10μl of OP50. Plates designated for stress observation were seeded with 10μl of OP50 and kept at room temperature for 5 minutes before use. Survival rates of L1 larvae on the plates were recorded at 24 and 48 hours, respectively.

### CRISPR/Cas9 gene editing and transgenic rescue

The CRISPR/Cas9 gene editing procedure was conducted following the methods described in previous literature ^32^. For detailed preparation see Section S2.

### Transcriptome sample preparation

Synchronized L1 larvae were used to transcriptome sampling. For detailed preparation see Section S3.

### Transcriptome sample sequencing

Total RNA was extracted using the Trizol (Invitrogen) reagent kit. RNA libraries were constructed using the NEB Next UltraTM RNA Library Prep Kit for Illumina (NEB, USA) with three biological replicates per treatment. The RNA libraries were sequenced using the Illumina NovaSeq 6000 platform.

After quality control, the clean reads were aligned to the reference genome using HISAT2 to obtain information about the reads’ mapping on the reference genome ^33^. Gene expression levels were quantified using feature Counts ^34^. Subsequently, differential expression analysis between contrast groups was conducted using DESeq2 ^35^. The gene expression levels were normalized using the Fragments Per Kilobase of transcript per Million mapped reads (FPKM) method. Principal component analysis (PCA) was performed on each sample using R. Significant DEGs which were subjected to Kyoto Encyclopedia of Genes and Genomes (KEGG) and Gene Ontology (GO) pathway enrichment analysis using KOBAS (KEGG Orthology Based Annotation System) ^36^. The statistical tests employed were hypergeometric distribution test and Fisher’s exact test, and p-values were adjusted using the Benjamini and Hochberg method.

### Quantitative Real-time PCR

The gene *tba-1* was utilized as an reference gene ^37^. Each biological sample, consisting of 500ng RNA, was reverse transcribed into cDNA using the TOYOBO ReverTra Ace R qPCR RT Master Mix with gDNA Remover kit (Catalog number: FSQ-301). The expression levels of each gene in the samples were validated using the Applied Biosystems QuantStudioTM6 Flex real-time PCR system and the TOYOBO SYBR Green detection system (Catalog number: QPK-201). Three biological replicates were performed for each treatment, with each biological replicate subjected to three technical replicates. Statistical analysis of the ΔCT values of each gene, normalized to the reference gene, under different treatment conditions was conducted using a two-tailed student’s t-test. The relative expression levels were calculated using the ΔΔCT method. For primers used see Table S3.

### Data analysis

Detailed methods for LOA, WormExp, and Chip-atlas analysis are described in Sections S4–S6.

### Microscopy

*hjSi224[vha-6p::DHS-3::GFP]* is a transgenic strain with overexpression of DHS-3::GFP fusion protein driving by *vha-6* promoter, DHS-3::GFP fusion protein is a well-established LD marker in *C. elegans* ^38–40^. *yqIs157[P_Y37A1B.5_mito-GFP]* is a transgenic strain with overexpression of mito::GFP fusion protein driving by *Y37A1B.5* promoter, mito::GFP fusion protein is a well-established mitochondrial marker in *C. elegans*^41^. For detailed methods see Section 7.

### Propidium Iodide (PI) staining

2.5 µl of 0.5 mg/mL PI (BioLegend, 421301) were added into 50 µl OP50 bacteria to prepare the food mixture^42^. For detailed methods see Section 8.

### RNA interference assay

RNAi constructs were generated by inserting target gene-specific sequences into the L4440 vector, which was then transformed into *Escherichia coli* HT115. *ftn-1* HT115 RNAi strain and *ads-1* HT115 RNAi strain were gifts from professor Dayong Wang ^43^. L4440 vector in HT115 feeding RNAi strain was used as a control for *gpx-1* and *gpx-7* RNAi assay. *gpx-1* OP50 RNAi strain and *gpx-7* OP50 RNAi strain were gifts from professor Lianfeng Wu ^44^. L4440 vector in OP50-based feeding RNAi strain was used as a control for *gpx-1* and *gpx-7* RNAi assay. For primers used see Table S3.

NGM agar plates supplemented with 1 mM IPTG and 50 ng/µL ampicillin were seeded with 200 µL of a 5x concentrated target RNAi bacterial for nematode cultivation ^45^. Three L4-stage larvae were transferred onto RNAi plates and incubated at 20°C. Gravid F1 adults were selected, and synchronized F2 L1 larvae were subsequently assessed for survival under 30‰ salinity conditions.

### MDA analysis

Tissue sample lysates were prepared through homogenization in liquid nitrogen ^43^, which was performed according to the protocol provided with the Malondialdehyde (MDA) Content Assay Kit (Micromethod, D799762-0100, Sangon). The MDA content was quantified by calculating the absorbance difference (ΔA532 - ΔA600) as specified in the kit instructions. To ensure the reliability and reproducibility of the results, three independent experiments were conducted.

### Statistical Analysis

Two-tailed student’s t-test was used in determination of significance between a pair of data. One-way ANOVA with Dunnett’s multiple comparisons test was used in determination of significance between groups in a trail with one time points. Two-way ANOVA with Dunnett’s multiple comparisons test was used in determination of significance between groups in a trial with two time points.

## Results

### *osm-11* mutants promote *C. elegans* salinity-stress tolerance

To answer which gene modulates *C. elegans* hyper-saline-stress tolerance, we examined 26 mutants (Figure 1A, B; Figure S1A, B), which were previously reported to be involved in osmoregulation under 30‰ salinity conditions. We found that *osm-11(rt142)* showed the strongest salinity-stress tolerance phenotype at both 24h and 48h after newly hatched L1 larvae were transferred to sea-water NGM plates (Figure 1A, B; Figure S1A, B).

**Figure 1.**
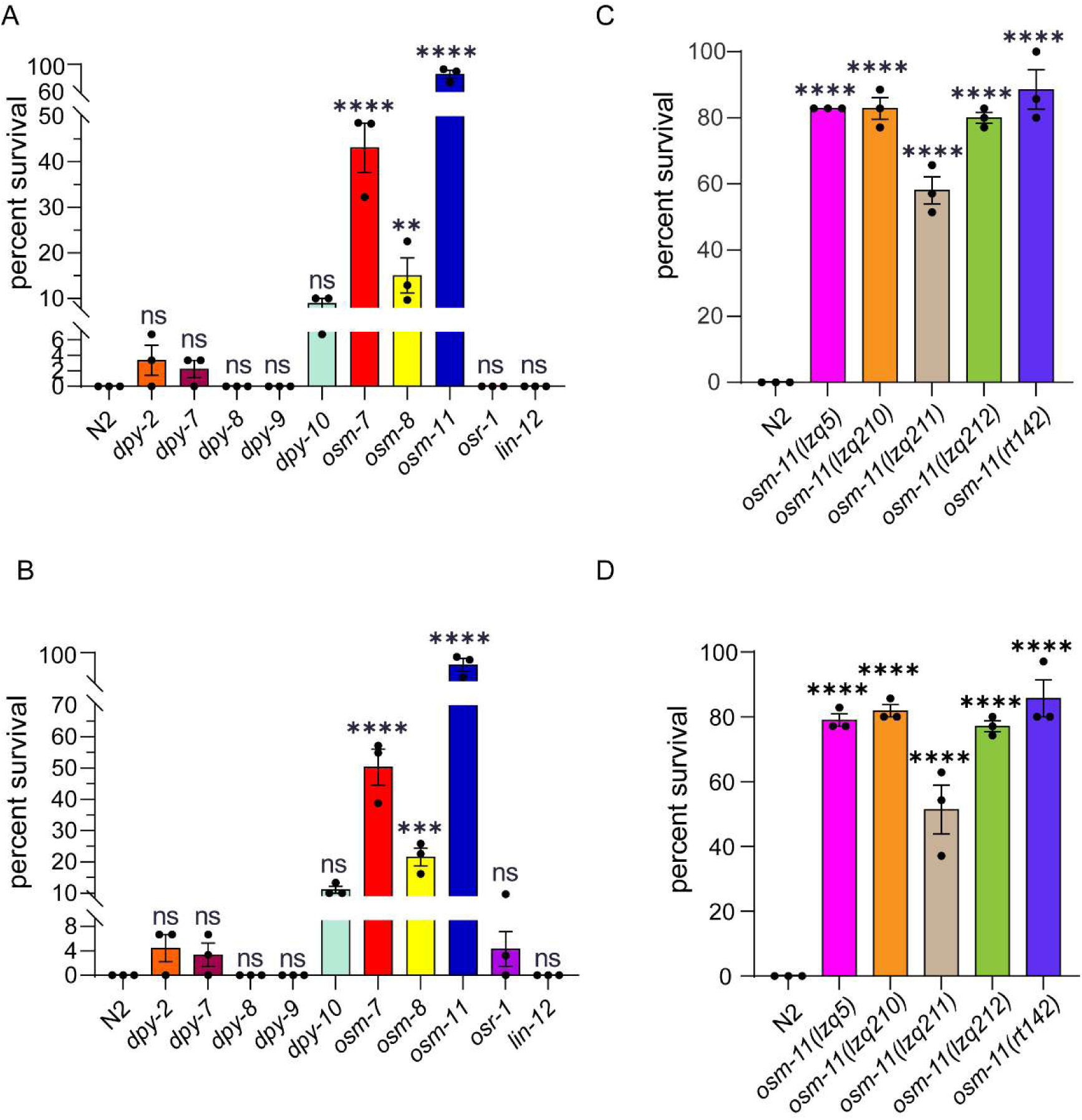
*osm-11* mutants promote *C. elegans* salinity-stress tolerance. (**A**) Survival rate of different mutants under 30‰ salinity environment at 24h. 30-35 synchronized L1 larvae were transferred to each plate, 3 replicates in each trial. (**B**) Survival rate of different mutants under 30‰ salinity environment at 48h. 30-35 L1 larvae in each replicate, 3 replicates in each trial. (**C**) Survival rate of *osm-11* mutants with different alleles under 30‰ salinity environment at 24h. 35 animals in each replicate, 3 replicates in each trial. (**D**) Survival rate of *osm-11* mutants with different alleles under 30‰ salinity condition at 48h. n = 35 animals in each replicate, 3 replicates in each trial. One-way ANOVA with Dunnett’s correction, Mean ± SEM. ****P-value < 0.0001, ***P-value < 0.001, **P-value < 0.01, *P-value < 0.05, ns = not significant.

Through CRISPR/Cas9 genome editing, we generated several *osm-11* mutants with different alleles (*lzq5*, *lzq210*, *lzq211*, and *lzq212*) (Figure S2A, B). All four mutants exhibited significantly enhanced hyper-saline tolerance compared to wild-type N2 worms (Figure 1C, D).

### Comparative transcriptome characterization between *osm-11* mutants and wild-type animals

To investigate how *osm-11* mutants promote *C. elegans* hyper-saline tolerance, we performed RNA sequencing on *osm-11(lzq5)* mutants and wild-type N2 animals under different salinity conditions (3‰, 6‰, 9‰, 12‰, 15‰ and 30‰) (Figure 2A; Figure S3A, B). Under a 30‰ hyper-saline environment, 1629 genes were significantly up-regulated, and 1192 genes were significantly down-regulated in *osm-11* mutants compared to wild-type animals (Figure S4F). Details to the significance of differentially expressed genes (DEGs) are presented in Data S1. Next, through KEGG enrichment analysis of the above up-regulated DEGs, we found that pathways such as fatty acid degradation, fatty acid biosynthesis, fatty acid metabolism and metabolism of xenobiotics by cytochrome P450 were significantly enriched (Figure 2B; Data S2). When KEGG analyses were performed on the down-regulated DEGs, we found that several pathways including calcium signaling were significantly enriched (Figure 2C; Data S3).

**Figure 2.**
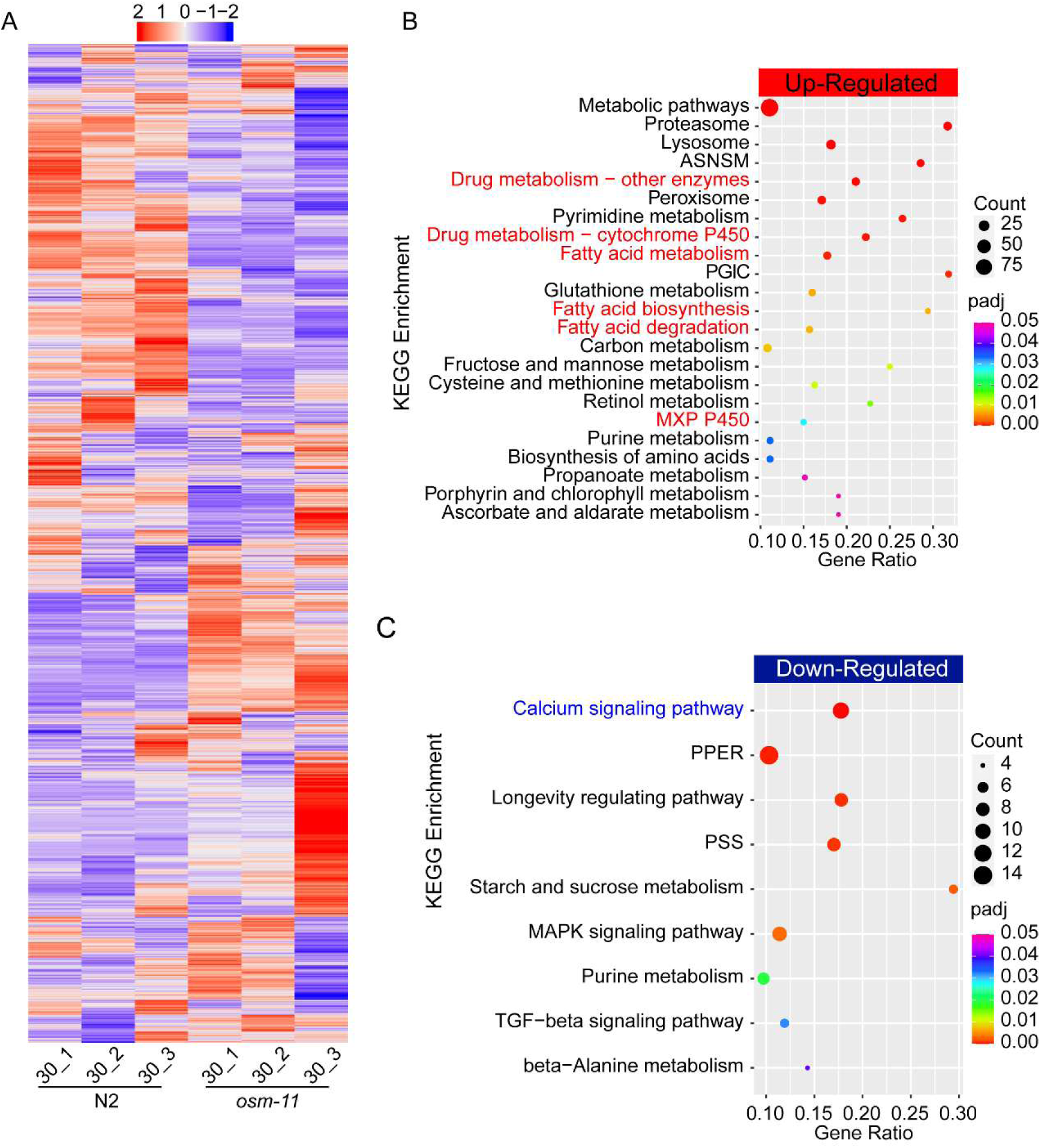
Transcriptome characterization between *osm-11* mutants and wild-type animals under 30‰ salinity condition. (**A**) Heatmap of DEGs (padj < 0.05 and fold change >1.5) under 30‰ salinity environment. (**B**) KEGG enrichment of up-regulated DEGs in *osm-11* mutants compared with wild-type animals. (**C**) KEGG enrichment of down-regulated DEGs in *osm-11* mutants compared with wild-type animals. ASNSM, Amino sugar and nucleotide sugar metabolism; PGIC, Pentose and glucuronate interconversions; MXC P450, Metabolism of xenobiotics by cytochrome P450; PPER, Protein processing in endoplasmic reticulum; PSS, Phosphatidylinositol signaling system.

Under other salinity conditions 3‰, 6‰, 9‰, 12‰, and 15‰ - 2849, 1169, 1629, 830 and 1579 genes were significantly up-regulated, respectively, while 2228, 808, 816, 389 and 883 genes were significantly down-regulated, respectively, in *osm-11* mutants compared to wild-type animals (Figure S4A-E). Details of the DEGs under each salinity condition were presented in Data S4. Next, through KEGG enrichment analysis of the above mentioned up-regulated DEGs, we found that pathways such as fatty acid metabolism, fatty acid degradation, fatty acid elongation, biosynthesis of unsaturated fatty acids and Drug metabolism - cytochrome P450 were significantly enriched in *osm-11(lzq5)* mutants in comparison to wild-type animals under most salinity conditions (Figure S3C; Data S5-9). For down-regulated DEGs, KEGG analysis showed that pathways including metabolism of xenobiotics by cytochrome P450 and Drug metabolism - cytochrome P450 were significantly enriched in *osm-11(lzq5)* mutants compared to wild-type animals under several salinity conditions (Figure S3D; Data S10-S14).

Interested genes from significantly enriched KEGG pathways including fatty acid metabolism, fatty acid degradation and biosynthesis of unsaturated fatty acids (Figure S5), were validated using quantitative Real-Time PCR (qPCR). Consistent trends between RNA-seq and qPCR are demonstrated in Figure S6A–E.

### *osm-11* mutants promoting salinity-stress tolerance depends on *acdh-12*

Given that fatty acid metabolism and P450 metabolism pathway-related genes were significantly increased in *osm-11* mutants compared with wild-type animals under most salinity conditions (Figure 2B; Figure S3C). We hypothesized that these genes up-regulated in *osm-11* mutants might contribute to their hyper-saline tolerance, so accordingly, knocking out these genes should reduce salinity-stress tolerance in *osm-11* mutants. To test whether knock out of up-regulated genes in *osm-11* mutants might reduce their salinity-stress tolerance, 11 up-regulated genes were selected to be knocked out in *osm-11* mutants. Most of the 11 up-regulated genes belong to KEGG pathways such as fatty acid metabolism and P450 metabolism, the expression of which were significantly increased in *osm-11* mutants compared with wild-type animals under most of salinity conditions (Figure S7). In addition, we observed that the expression of *acdh-12*, *acs-17*, *ugt-18* and *fat-6* was significantly increased in wild-type N2 worms when salinity conditions changed from 15‰ to 30‰ (Figure S8C). Similar to wild type animals, we found that the expression of *acdh-12*, *ech-9*, *acs-17*, *ugt-18* and *fat-6* were significantly up-regulated in *osm-11* mutants when salinity increased from 15‰ to 30‰ (Figure S8C). Through CRISPR/Cas-9 genome editing, we successfully generated 33 mutants in 9 genes out of the 11 up-regulated genes in wild-type N2 background (Data S15). Then we generated double mutants by crossing single mutants with *osm-11(lzq5)* mutants. From this we observed that o*sm-11;acdh-12*, *osm-11;acs-17* and *osm-11;ugt-15* double mutants showed significantly reduced survival rate in comparison to *osm-11* single mutants, suggesting that salinity-stress tolerance in *osm-11* mutant depends on *acdh-12*, *acs-17* and *ugt-15* (Figure 3A, B). *acdh-12*, *acs-17* and *ugt-15* encode Acyl CoA Dehydrogenase 12, fatty Acid CoA Synthetase and UDP-Glucuronosyl Transferase 15, respectively. *acdh-12* and *acs-17* are involved in fatty acid metabolism, while *ugt-15* might function in cytochrome p450 metabolism.

**Figure 3.**
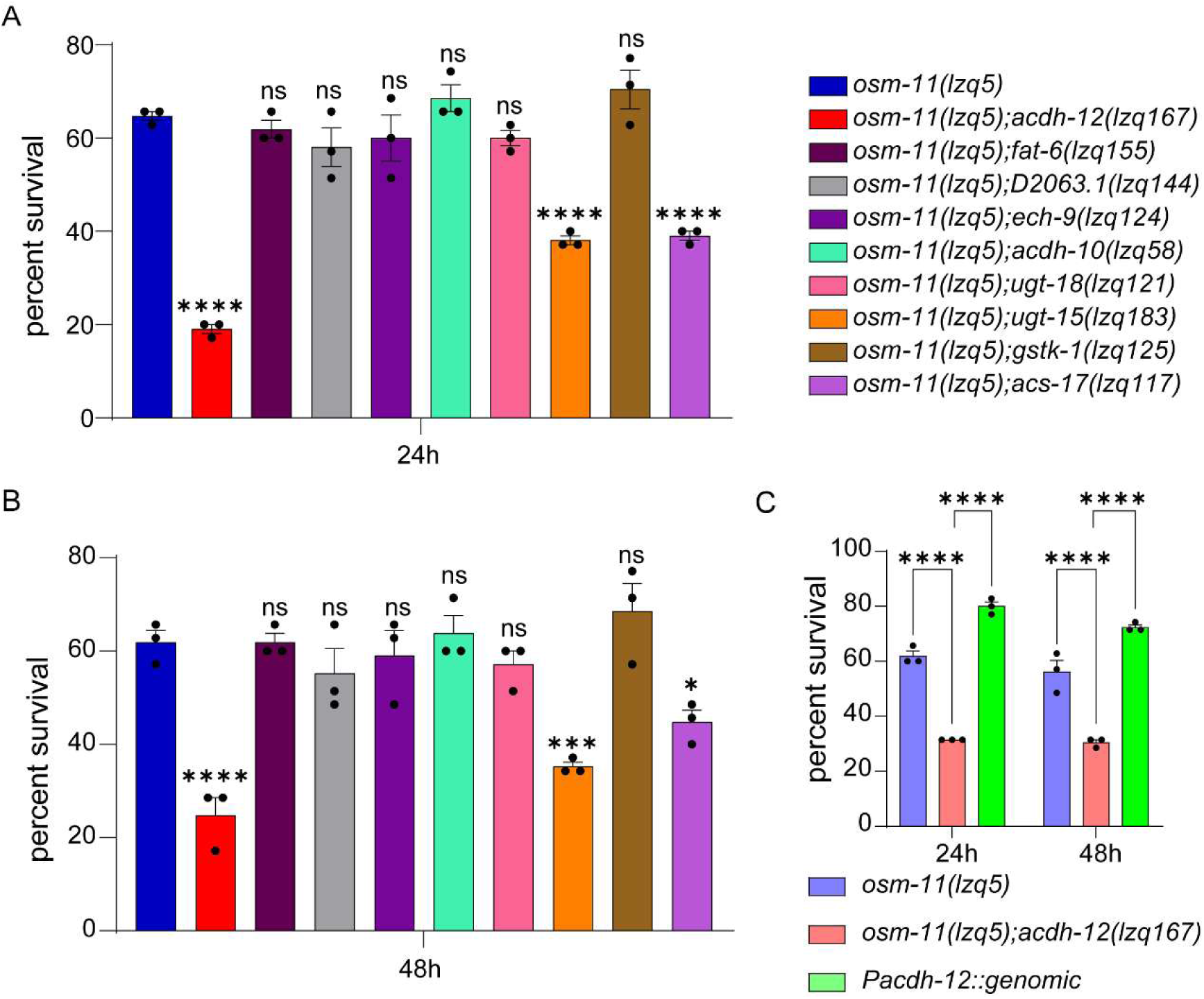
Mutation in *osm-11* promoting salinity-stress tolerance depend on *acdh-12*. (**A**) Survival rate of double mutants at 24h under hyper-saline condition. (**B**) Survival rate of double mutants under hyper-saline condition at 48h. (**C**) Overexpression of wild-type *acdh-12* genomic DNA recused the reduced survival rate of *osm-11;acdh-12* double mutants under hyper-saline conditions at 24h and 48h, respectively. n = 35 animals in each replicate, 3 replicates in each trial. Two-way ANOVA with Dunnett’s correction, Mean ± SEM. *P-value < 0.05, **P-value < 0.01, ***P-value < 0.001, ****P-value < 0.0001, ns, not significant.

Given that, mutation in *acdh-12* reduced the survival rate more severely in *osm-11* mutants under hyper-saline condition, compared to mutations in either *acs-17* or *ugt-15* (Figure 3A, B). Next, we focused our study on how mutation in *acdh-12* impaired salinity-stress tolerance in *osm-11* mutants. Through overexpression of the wild-type *acdh-12* genomic DNA driving by *acdh-12* promoter, we rescued hyper-saline resistance in *osm-11;acdh-12* double mutants to the levels of *osm-11* single mutants (Figure 3C). In aggregate, this data indicates that *acdh-12* is required for hyper-saline tolerance in *osm-11* mutant.

We hypothesized that genes down-regulated in *osm-11* mutants might suppress salinity-stress tolerance and knock out these genes in wild-type worms, which would increase hyper-saline tolerance. To test whether knock out of the down-regulated genes in wild-type animals might increase their hyper-saline resistance, 5 down-regulated genes belonging to calcium signaling were selected. The expression of these 5 calcium signaling genes were significantly decreased in *osm-11* mutants compared with wild-type animals under most salinity conditions (Figure S7). Through CRISPR/Cas9 genome editing, we successfully generated 16 mutants in 4 out of 5 genes in N2 background (Data S15). We found that mutations in either *gar-3* or *ckr-2* promoted salinity tolerance under 25‰, but not 30‰ salinity condition, while mutation in *ncx-1* reduced the tolerance under 25‰ condition, but not 30‰ (Figure S9).

Furthermore, we generated triple mutants by crossing *osm-11;acdh-12* double mutants with other double mutants generated in this study (Figure 3A, B), to explore whether *acdh-12* functions in the same pathway or in parallel with other candidate genes (Figure S10). Unexpectedly, we observed that *osm-11;acdh-12;ugt-15*, and *osm-11;acdh-12;acs-17* triple mutants exhibited higher survival rate compared to *osm-11;acdh-12* double mutants (Figure S10).

### Comparative transcriptome characterization between *osm-11* single mutants and *osm-11;acdh-12* double mutants

To answer how mutation in *acdh-12* reduces salinity-stress tolerance in *osm-11* single mutants, we applied RNA-seq on *osm-11* single mutants and *osm-11;acdh-12* double mutants under 3‰ and 30‰ salinity conditions, respectively (Figure 4A). Principal component analysis (PCA) showed that the samples are separated according to different salinity conditions and genetic background (Figure 4B). Under 30‰ hyper-saline condition, 2960 genes were significantly up-regulated, and 2811 genes were significantly down-regulated in *osm-11* single mutants compared to *osm-11;acdh-12* double mutants (Figures S11A). Details of the significant DEGs were presented in Data S16. Next, through KEGG analysis of these up-regulated DEGs, we found that pathways such as oxidative phosphorylation, metabolism of xenobiotics by cytochrome P450, fatty acid metabolism, and the citrate cycle (TCA cycle) were significantly enriched in *osm-11* mutants compared with *osm-11;acdh-12* double mutants (Figure 4C, Data S17). When KEGG analysis was applied to the down-regulated DEGs, pathways including autophagy, calcium signaling, and mitophagy there was significant enrichment in *osm-11* single mutants in comparison to *osm-11;acdh-12* double mutants (Figure S11B; Data S18). Furthermore, we found that the expression of genes in many mitochondria related GO terms were significantly increased in *osm-11* mutants compared to *osm-11;acdh-12* double mutants (Data S19). This data suggested that mutation in *acdh-12* might reduce salinity-stress tolerance of *osm-11* mutants via up-regulation of genes in calcium signaling, autophagy and mitophagy pathways, and downregulation of genes in pathways such as oxidative phosphorylation and the TCA cycle.

**Figure 4.**
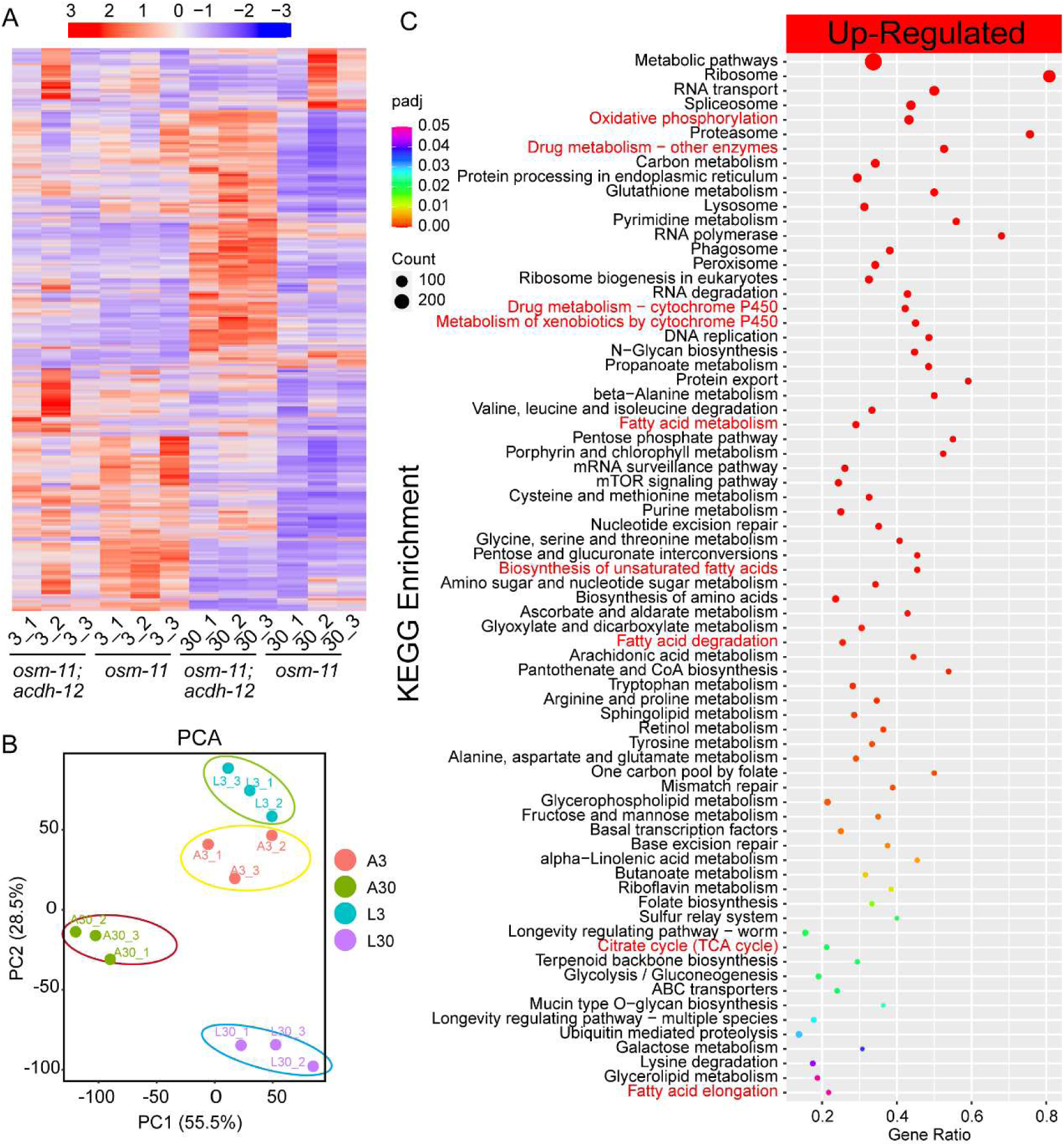
Transcriptome characterization between *osm-11* single mutants and *osm-11;acdh-12* double mutants. (**A**) Heatmap of DEGs (padj < 0.05, foldchange > 1.5) under 3‰ and 30‰ salinity environment (respectively. (**B**) PCA of *osm-11(lzq5)* and *osm-11;acdh-12* transcriptomes. A represents *osm-11;acdh-12* mutant, L represents *osm-11(lzq5)* mutant. (**C**) KEGG enrichment of up-regulated DEGs in *osm-11(lzq5)* vs. *osm-11;acdh-12* under 30‰ salinity.

### acdh-12 might regulate salinity-stress tolerance of *osm-11* mutants via *dhs-28* and *daf-22*

*acdh-12* is reported to be involved in the assembly of respiratory complex I in the mitochondria ^46^, although it was also proposed to be essential to fatty acid oxidation. We noted that the expression of genes such as *dhs-28* and *daf-22* in KEGG pathways “biosynthesis of unsaturated fatty acids”, were significantly up-regulated in *osm-11* mutants compared with *osm-11;acdh-12* double mutants (Figure 4C; Figure S12B, D). Interestingly, we also found that *dhs-28* was significantly increased in *osm-11* mutants compared with wild-type N2 animals under 30‰ salinity condition, and *daf-22* was increased in *osm-11* mutants compared to N2, although without statistics significance (Figure S12A, C). Then, we crossed *dhs-28* or *daf-22* single mutants ^32^ with *osm-11* mutants, to generate *osm-11*;*dhs-28* and *osm-11;daf-22* double mutants, respectively. Notably, we observed that both double mutants showed significance in reduced survival rate compared with *osm-11* single mutants, under 30‰ salinity conditions (Figure S12E), indicating that salinity-stress tolerance of *osm-11* mutants depended on *dhs-28* and *daf-22*, which were positively modulated by *acdh-12*.

Given that both *dhs-28* and *daf-22* are involved in peroxisomal β-oxidation relating to lipid catabolism and homeostasis, we asked if the number and size of lipid droplets were changed in *osm-11* mutants compared with *osm-11;acdh-12* double mutants or wild-type animals (Figure S13A). Through examining DHS-3::GFP labeled lipid droplets (LDs) in the worms’ intestine, we observed a significant reduction in the number of lipid droplets (LDs) in both wild-type N2 and *osm-11* mutants after a 3 hour treatment under 30‰ salinity conditions, compared to 3‰ salinity. By contrast, the number of lipid droplets in *osm-11;acdh-12* double mutants increased significantly following 30‰ salinity treatment (Figure S13B). It is also worth noting that we observed that the size of LDs was significantly increased in wild-type N2 animals. However, it was significantly reduced in *osm-11* mutants when animals were under salinity stress, while there was no significant size change in *osm-11;acdh-12* double mutants after being subjected to hyper-saline stress (Figure S13C). Mutations in either *dhs-28* or *daf-22* were reported to induce enlarged LDs ^47^. Our data suggested that salinity-stress tolerance in *osm-11* mutants may be contributed to the up-regulation of *dhs-28* and *daf-22*, resulting in a reduction in lipid droplet size.

### *acdh-12* dependent and independent transcriptome characterizations in response to hyper-saline environment

To investigate whether transcriptome characterizations in *osm-11* mutants depend on *acdh-12*, we applied Luperchio Overlap Analysis (LOA) based on Vora’s method ^48^ to obtain the overlapping and pair-specific DEGs between *osm-11* single mutants vs. N2 and *osm-11* single mutants vs. *osm-11;acdh-12* double mutants under 30‰ salinity conditions. A total of 1672 overlapping DEGs were found between *osm-11* vs. N2 and *osm-11* vs. *osm-11;acdh-12* (Figure S14A; Data S21). However, 251 of the overlapping DEGs were not positively correlated (e.g., a gene is up-regulated in *osm-11* vs. N2 but down-regulated in *osm-11* vs. *osm-11;acdh-12*), suggesting that these genes might be *acdh-12* dependent (Figure S14B; Data S21). Accordingly, there were 1421 positively correlated overlapping DEGs acting in an *acdh-12* independent manner (Figure S14B; Data S21). Among the 1421 positively correlated overlapping DEGs, 786 up-regulated overlapping ones were enriched in KEGG pathways - such as phagosome, metabolism of xenobiotics by cytochrome P450, drug metabolism-other enzymes, proteasome, peroxisome, glutathione metabolism, and fatty acid degradation (Figure S14C; Data S21) - while 635 down-regulated overlapping genes were enriched in KEGG pathways including calcium signaling pathway, spliceosome and longevity regulating pathway-multiple species (Figure S14D; Data S21). From these 251 non-positively correlated DEGs, enrichment was observed in KEGG pathways including the metabolism of xenobiotics by cytochrome P450 and autophagy – animal (Figure S14E; Data S21).

Furthermore, 2043 DEGs specific to *osm-11* vs. N2 and 3888 DEGs specific to *osm-11* vs. *osm-11;acdh-12* might be *acdh-12* dependent and depended on mutation in *acdh-12*, respectively (Figure S14B). For up-regulated DEGs specific to *osm-11* vs. N2, KEGG pathways including metabolic pathways, ABC transporters and Endocytosis were significantly enriched (Figure 5A; Data S22), while KEGG pathways such as TGF-beta signaling pathway, Wnt signaling pathway and Ubiquitin mediated proteolysis, were significantly enriched in down-regulated DEGs specific to *osm-11* vs. N2 (Figure 5B; Data S22). In up-regulated DEGs specific to *osm-11* vs. *osm-11; acdh-12*, the most significantly enriched KEEG pathways included Spliceosome, Oxidative phosphorylation, Peroxisome, Lysosome and Drug metabolism-other enzymes (Figure 5C; Data S22). Oxidative phosphorylation was a prominent KEGG pathway enriched in the up-regulated DEGs specific to *osm-11* vs. *osm-11; acdh-12* (43/111, Data S22), indicating that reduced Oxidative phosphorylation in *osm-11;acdh-12* double mutants might lead to impaired energy supply, which might confer their reduced salinity-stress tolerance.

**Figure 5.**
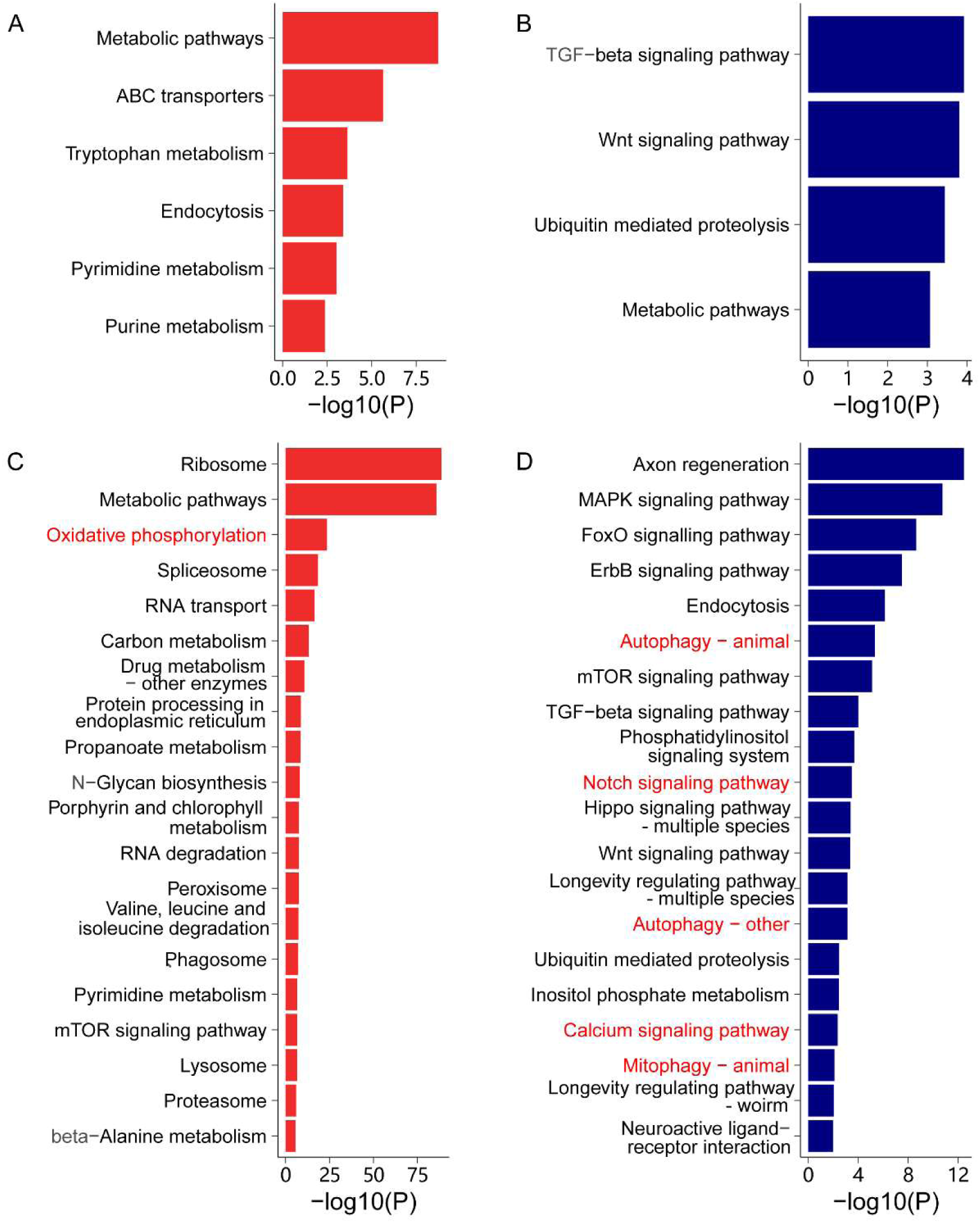
Assessing the non-overlapped KEGG pathways between *osm-11* vs. *N2* and *osm-11* vs. *osm-11; acdh-12*. (**A**) KEGG enrichment of up-regulated DEGs specific to *osm-11* vs. N2. (**B**) KEGG enrichment of up-regulated DEGs specific to *osm-11* vs. N2. (**C**) KEGG enrichment of up-regulated DEGs specific to *osm-11* vs. *osm-11;acdh-12*. (**D**) KEGG enrichment of down-regulated DEGs specific to *osm-11* vs. *osm-11;acdh-12*. All KEGG pathways were filtered for a statistical threshold of q < 0.05. Only the top 20 terms are shown.

Similarly, GO terms such as “mitochondrion” (GO:0005739, 193/525), “mitochondrial large ribosomal subunit” (GO:0005763, 23/27; GO:0005762, 30/34), “mitochondrial small ribosomal subunit” (GO:0022627, 30/32), “mitochondrial translation”(GO:0032543, 26/31) and “oxidation-reduction process” (GO:0055114, 107/539), were significantly enriched in the up-regulated DEGs specific to *osm-11* vs. *osm-11; acdh-12* (Data S22). In contrast, no GO terms associated with mitochondrion were found in up-regulated DEGs specific to *osm-11* vs. *N2* (Data S22). These results indicate that *osm-11;acdh-12* might have much reduced mitochondrial synthesis and function, as well as reduced energy supply and oxidation-reduction ability compared to *osm-11* single mutants, which might lead to reduced salinity-stress tolerance in *osm-11;acdh-12* double mutants compared to *osm-11* single mutants.

In down-regulated DEGs specific to *osm-11* vs. *osm-11;acdh-12*, the most significantly enriched KEGG pathways contained Autophagy (cel04140, cel04136), Mitophagy (cel04137) and Fatty acid metabolism (Figure 5D; Data S22). The down-regulated genes in Autophagy pathway (cel04140, cel04136) included *daf-15, atg-13, mom-4, aak-1, rsks-1, pdk-1, atg-16.1, atg-18, ist-1, pek-1, vamp-8, atg-2, cpl-1, atg-9, snap-29*, while genes in mitophagy pathway (cel04137) included *daf-16*, *src-1*, *pek-1*, *atg-9* and *tbc-14* (Data S22), indicating that enhanced autophagy and mitophagy in *osm-11;acdh-12* double mutants might contribute to their decreased survival rate compared to *osm-11* single mutants under hyper-saline environment.

### Mitochondrial fragmentation might contribute to hyper-saline sensitivity

As we discovered that the expression of genes in mitochondrial function related pathways, such as oxidative phosphorylation and TCA cycle, was significantly repressed in *osm-11;acdh-12* double mutants compared to *osm-11* single mutants under hyper-saline environment (Figure 4C and 5C; Data S17; Data S19), given that mitochondrial functions are always associated with mitochondria morphology changes, we then asked whether mitochondria morphology changed in hyper-saline conditions and tested it by crossing a florescence labeled strain *yqIs157 (P_Y37A1B.5_mito-GFP)* ^49^, which was specially expressed in the mitochondria of hypodermis, to both *osm-11* single mutants and *osm-11;acdh-12* double mutants. From this, we observed that most L4 wild-type N2 worms exhibited mitochondrial fragmentation after 3h exposure to 30‰ salinity, while both *osm-11* single mutants and *osm-11;acdh-12* double mutants showed much less mitochondrial fragmentation compared to wild-type animals (Figure 6A, B). In addition, mitochondrial fragmentation at 6h, 12h, 24h and 48h under 30‰ salinity conditions was prevalent in wild-type L4 worms but not in *osm-11* single mutants or *osm-11;acdh-12* double mutants at 6h, 12h, 24h or 48h under 30‰ salinity conditions (Figure 6B). Furthermore, we found that mitochondrial fragmentation at 3h, 6h, 12h and 24h under 30‰ salinity conditions was also prevalent in wild-type L1 worms but not in *osm-11* single mutants or *osm-11;acdh-12* double mutants (Figure 6C). Moreover, we observed that the percentage of intermediate mitochondrial fragmentation was significantly increased in the L1 larvae of *osm-11;acdh-12* double mutants in comparison to *osm-11* single mutants at 12h and 24h after subjected to 30‰ salinity conditions (Figure 6C). Collectively, our data suggested that mitochondrial fragmentation might contribute to hyper-saline sensitivity in wild-type animals, while mutation in *osm-11* reduced mitochondrial fragmentation, and *acdh-12* dependent mitochondrial fragmentation regulation was developmental stage specific. This may contribute to the reduced salinity-stress tolerance observed in *osm-11;acdh-12* double mutants compared to *osm-11* single mutants.

**Figure 6.**
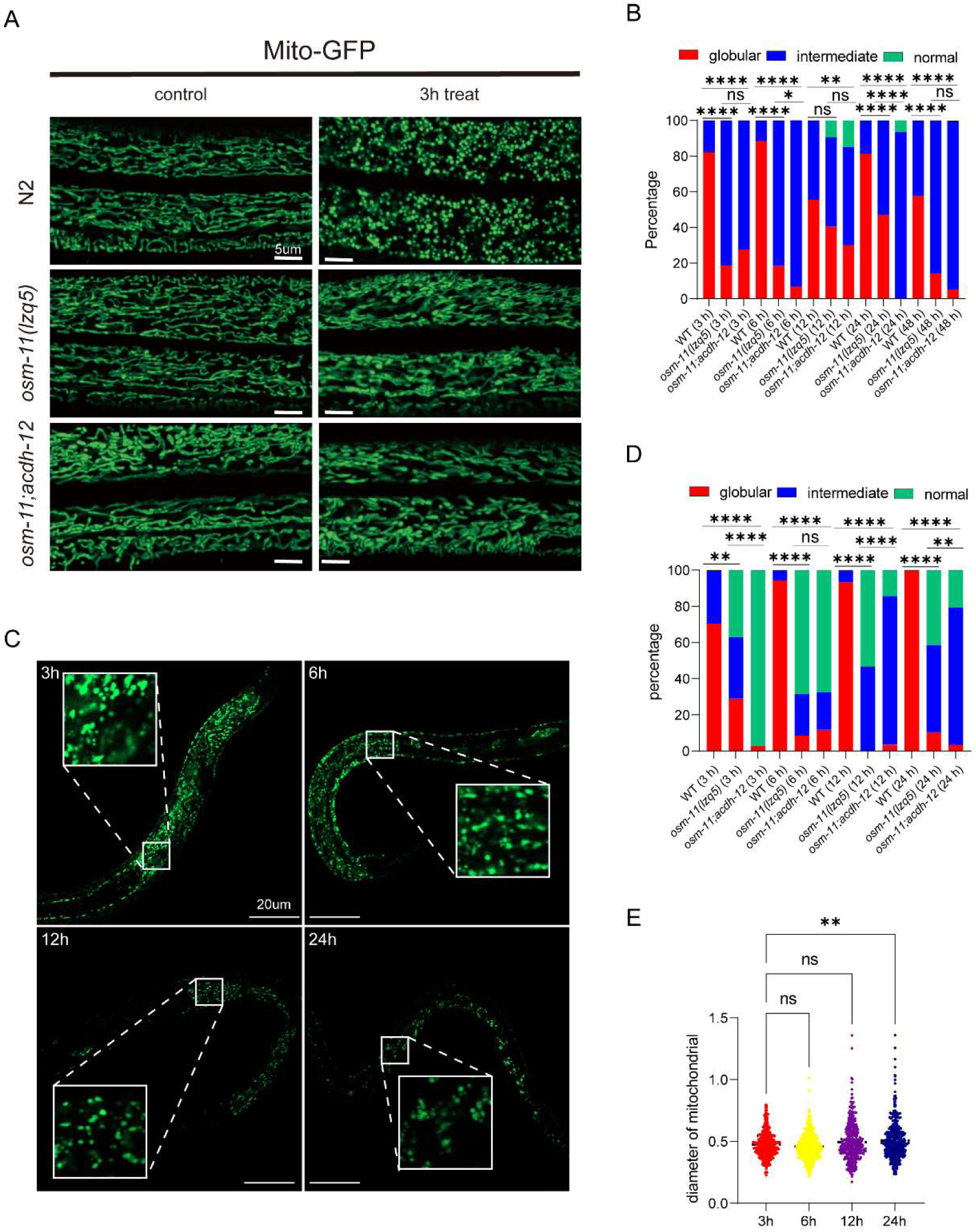
Mitochondrial fragmentation might contribute to hyper-saline sensitivity. (**A**) Representative images of mitochondria in the hypodermis of wild-type N2, *osm-11* single mutants and *osm-11;acdh-12* double mutants, L4 larvae were used under 30‰ salinity and control conditions. Bar, 5μm; n ≥ 30 individuals were scored for each trial. (**B**) Percentage of different mitochondrial morphology under 30‰ salinity at different time points. L4 larvae were used for imaging. n ≥ 30 individuals were scored for each trial. *P-value < 0.05, **P-value < 0.01, ****P-value < 0.0001, ns, not significant, Fisher’s exact test. (**C**) Percentage of different mitochondrial morphology under 30‰ salinity at different time points. L1 larvae were used for imaging. n ≥ 30 individuals were scored for each trial. **P-value < 0.01, ****P-value < 0.0001 (Fisher’s exact test. (**D**) Representative images of mitochondria in the hypodermis of wild-type animals at different time points, L1 larvae were applied for imaging under 30‰ salinity conditions. Bar, 20μm; n ≥ 35 individuals was scored for each trial. (**E**) The size of fragmented mitochondria under 30‰ salinity at different time points, L1 larvae were applied for imaging. Each point represents one mitochondrion. n = 366, 576, 365 and 445 for 3h, 6h, 12h and 24h, respectively. Over 20 individuals were scored in each time point. **P-value < 0.01, One-way ANOVA with Dunnett’s correction.

It is important to note that all wild-type L1 larvae were dead after 24 hours under 30‰ salinity conditions (Figure 1A, B), and mitochondria fragmentation was prevalent at all time points (Figure 6C). We then asked if the size of fragmented mitochondria change from 3h to 24h. Interestingly, we observed that the size of fragmented mitochondria had significantly increased at 24h compared to other time points (Figure 6D, E), suggesting that the enlarged size of fragmented mitochondria might confer animal death under salinity-stress environment.

To study the mechanism by which mitochondrial fragmentation confers animal death in hyper-saline conditions, we examined cell and membrane integrity via propidium iodide (PI) staining. Compared to wild-type animals, when L4 larvae were transferred to hyper-saline environments, we observed that *osm-11* single mutants as well as *osm-11;acdh-12* double mutants exhibited significantly high survival rates at both 24h and 48h (Figure 7A). Meanwhile, the mean fluorescence intensity of wild-type animals was significantly increased compared to both *osm-11* single mutants and *osm-11;acdh-12* double mutants (Figure 7*B*). Collectively, this data suggested that mitochondrial fragmentation was positively correlated with the impairment of cell and membrane integrity, and mutation in *osm-11* blocked mitochondrial fragmentation and maintained cell and membrane integrity to increase salinity-stress tolerance.

**Figure 7.**
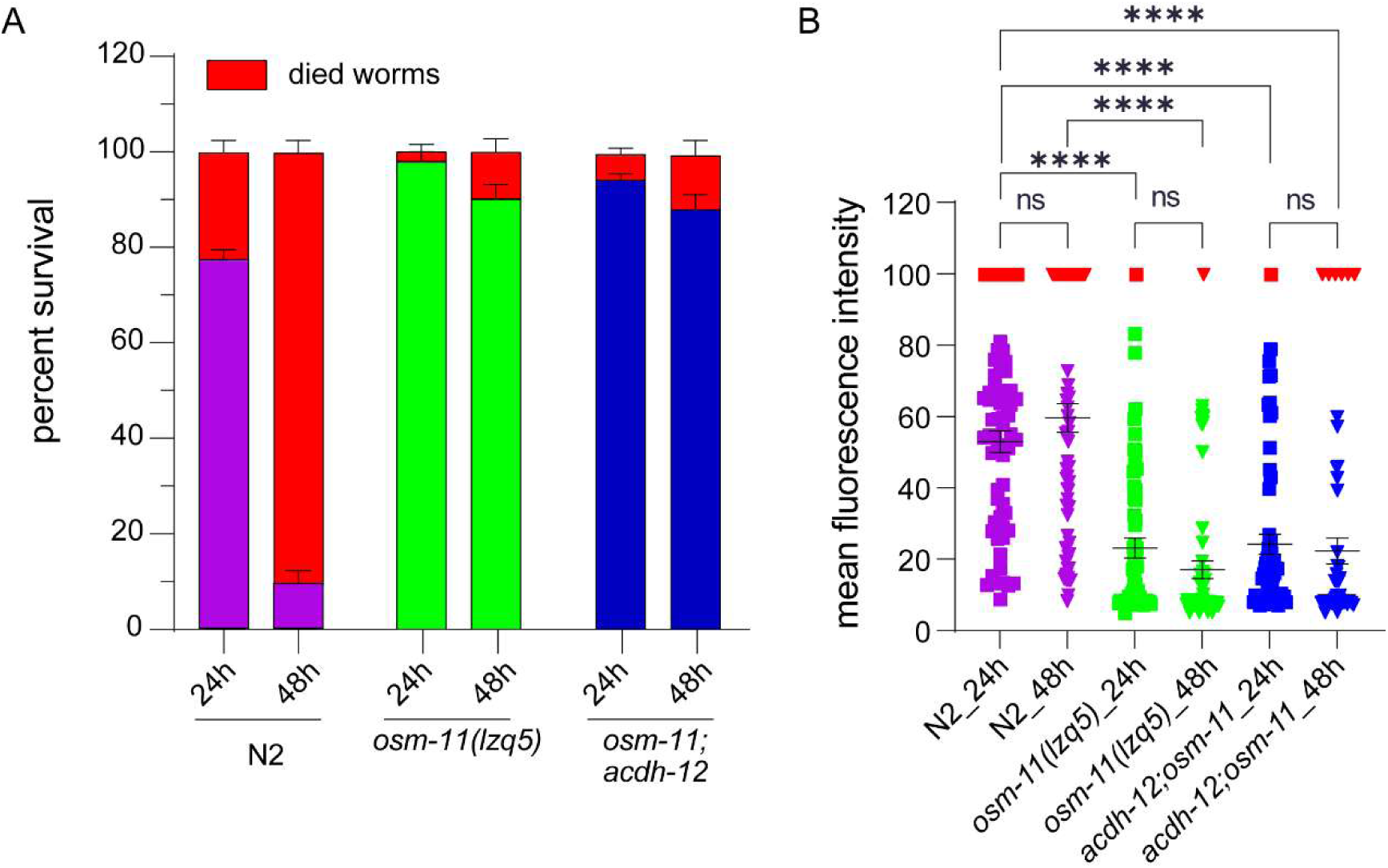
Cell and membrane integrity might contribute to hyper-saline resistance. (**A**) Survival rate of L4 larvae of N2, *osm-11(lzq5)* and *osm-11;acdh-12* under 30‰ salinity at 24h and 48h. Three trials, n = 35 worms on average in each trial. (**B**) Intensity of PI staining for wild-type N2, *osm-11* single mutants and *osm-11;acdh-12* double mutants, under 30‰ salinity at 24h and 48h. Each dot represents one individual. ****p < 0.0001, ns, not significant, Mann-Whitney with non-parameter test. Red color labeled icons represent died worms.

### *osm-11* mutants might promote salinity-stress tolerance via ferroptosis inhibition

Given that ferroptosis inhibition related genes such as *ftn-1* were significantly up-regulated in *osm-11* mutants compared with wild-type animals (Data S1), and both *ftn-1* and *ftn-2* were significantly down-regulated in *osm-11;acdh-12* double mutants in comparison to *osm-11* single mutants (Data S16), we asked whether the salinity-stress tolerance of *osm-11* mutants contributed by ferroptosis inhibition Dihomo-γ-Linolenic Acid (DGLA) was reported to trigger ferroptosis in the germ cells and sterility in *C. elegans* ^50^. Through DGLA supplementation to both *osm-11* mutants and wild-type animals, we found that DGLA supplementation significantly reduced salinity-stress tolerance in *osm-11* mutants (Figure S17), indicating that ferroptosis inhibition might contribute to hyper-saline resistance.

Furthermore, we found that in addition to *ftn-1* and *ftn-2* (which encode ferritin), the expression of *ads-1* (which encodes alkylglycerone phosphate synthase), *gpx-1* and *gpx-7* (which encodes glutathione peroxidases GPX4), *dhod-1* (which encodes dihydroorotate dehydrogenase), as well as *mboa-6* and *mboa-7* (which encodes phospholipid-modifying enzymes/acetyltransferases) were significantly down-regulated in *osm-11;acdh-12* double mutants in comparison to *osm-11* single mutants (Data S16). All the above-mentioned genes are known to inhibit ferroptosis, suggesting that the *acdh-12* mutation might promote ferroptosis via downregulation of ferroptosis protective genes, leading to reduced salinity-stress tolerance in *osm-11* mutants.

To test if these ferroptosis protective genes are necessary to promote animal salinity-stress tolerance, we knocked down the genes via RNA interference (RNAi) in *osm-11* mutants. Intriguingly, we observed that simultaneous knockdown of both *gpx-1* and *gpx-7* genes significantly reduced the survival rate of *osm-11* mutants, under 30‰ salinity environment (Figure 8A, B), while no significant decrease in survival rate was observed following RNAi knockdown of *ftn-1, ads-1*, *dhod-1,* and *mboa-6*, respectively (Figures S18). Thus, our data suggested that ferroptosis protective genes *gpx-1* and *gpx-7* contributed to the salinity-stress tolerance of *osm-11* mutants, and mutation in *acdh-12* might lead to ferroptosis by modulating both *gpx-1* and *gpx-7* at the transcriptional level.

**Figure 8.**
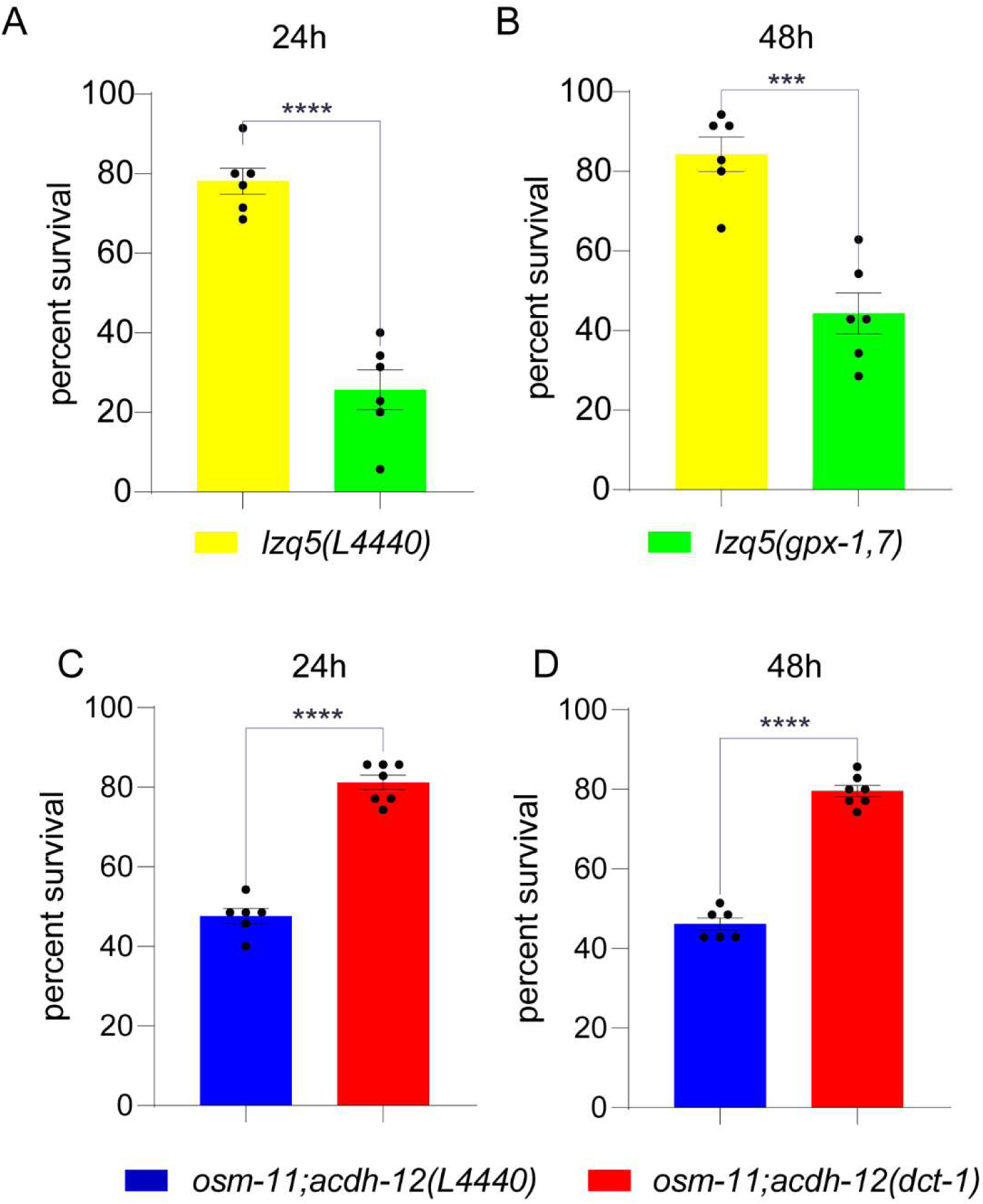
Genes involved in ferroptosis and mitophagy contribute to salinity-stress tolerance in *osm-11* mutants. (**A**) Survival rate of *osm-11* mutants was significantly reduced by *gpx-1/gpx-7* RNAi at 24h, under 30‰ salinity environment. *osm-11* treated with L4440 (OP50) was used as the control. (**B**) Survival rate of *osm-11* mutants was significantly reduced by *gpx-1/gpx-7* RNAi at 48h, under 30 ‰ salinity environment. *osm-11* treated with L4440 (OP50) was used as the control. (**C**) Survival rate of *osm-11;acdh-12* double mutants was significantly increased by *dct-1* RNAi at 24h, under 30‰ salinity environment. *osm-11;acdh-12* treated with L4440 (HT115) was used as the control. (**D**) Survival rate of *osm-11;acdh-12* double mutants was significantly increased by *dct-1* RNAi at 48h, under 30‰ salinity environment. *osm-11;acdh-12* treated with L4440 (HT115) was used as the control. 35 synchronized L1 larvae were transferred to each plate. Student’s t-test, Mean ± SEM. ***P-value < 0.001, ****P-value < 0.0001.

The level of Malondialdehyde (MDA) was reported to be increased during ferroptosis and an indicator of lipid peroxidation ^51^. From this information, we then measured the malondialdehyde (MDA) levels in both wild-type animals and *osm-11* mutants, under normal physiological conditions, and found that the level of MDA in *osm-11* mutants was significantly lower than that in wild-type animals (Figure S19), suggesting that mutation in *acdh-12* might reduce salinity-stress tolerance of *osm-11* mutants via increased lipid peroxidation and subsequent ferroptosis.

### *acdh-12* mutation might reduce salinity-stress tolerance via mitophagy

Given that the expression of mitophagy receptor gene *dct-1* was significantly up-regulated in *osm-11;acdh-12* double mutants in comparison to *osm-11* single mutants (Data S16), we performed RNA interference (RNAi) knockdown of *dct-1* in *osm-11;acdh-12* double mutants to test if the decreased survival rate of double mutants was due to mitophagy. We found that *dct-1* knockdown significantly increased the survival rate of *osm-11;acdh-12* double mutants, under 30‰ salinity condition (Figure 8C, D; Figure S20). To further confirm our *dct-1* RNAi results, we designed a new RNAi plasmid, *dct-1* (B) RNAi, and as expected, *dct-1* (B) RNAi also significantly promoted salinity-stress tolerance in *osm-11;acdh-12* double mutants (Figure 1). Furthermore, we observed that *daf-16*, *pha-4*, and *hlh-30*, being transcription factors regulating *dct-1* expression, were also significantly up-regulated in *osm-11;acdh-12* double mutants in comparison to *osm-11* single mutants, under salinity-stress condition (Data S16). Together, our data indicated that *acdh-12* mutation might reduce salinity-stress tolerance via mitophagy, in line with significantly repressed expression of genes in TCA cycle and oxidative phosphorylation pathways (Data S17- overall showing that mitophagy and impaired function of mitochondrial TCA cycle and oxidative phosphorylation could induce energy crisis and increased cell death in *osm-11;acdh-12* double mutants.

## Discussion

Salinization is an overlooked consequence of global climate change, which is currently occurring at an unprecedented rate. Salinization increasingly threatens terrestrial soil and freshwater organisms, as it can induce tremendous physiological costs and eventually provoke mortality ^52^. Given that most terrestrial animals tolerate only limited salinity ranges, it is essential to understand how these hyposaline-sensitive animals could adapt to the threat of aggressive salinity stress. In this research, we investigate the mechanism underlying how mutation in Notch co-ligand encoding gene *osm-11* promotes salinity-stress tolerance. Using candidate mutants screening and CRISPR/Cas-9 genome editing, we found that, to our knowledge, *osm-11* mutants exhibited the strongest salinity-stress tolerance phenotype in *C. elegans*. Next, through comparative transcriptomics and genome editing, we found that salinity-stress tolerance of *osm-11* mutants depended on *acdh-12* encoding an acyl-CoA dehydrogenase, which might contribute to lipid β-oxidation in mitochondria ^53^. Recently, it was reported that *acdh-12* functions as an assembly factor for mitochondrial complex I, which are essential for mitochondrial respiration and energy supply ^46^. We found that mutation in *acdh-12* significantly reduced the survival rate of *osm-11* mutants under salinity stress conditions, indicating that salinity-stress tolerance of *osm-11* mutants depend on *acdh-12*.

Mitochondria play a crucial role in maintaining cellular health, with functions include, but are not limited to, ATP synthesis, lipid metabolism, heme biosynthesis, as well as the regulation of cellular metabolism, stress responses, and cellular fate ^54–56^. In this study, we observed that under hyper-saline conditions, compared to *osm-11* single mutants, the expression of genes related to oxidative phosphorylation was significantly repressed in *osm-11;acdh-12* double mutants, while the expression of genes related to autophagy and mitochondrial autophagy pathways was significantly up-regulated (Figure S11B). The significantly down-regulated genes in the oxidative phosphorylation pathway included: *asb-1*, *asg-1*, *asg-2*, *atp-3*, *atp-5*, *C14B9.10*, *C16A3.5*, *cox-5A*, *cox-5B*, *cox-6A*, *cox-6B*, *cox-6C*, *cox-7C*, *cox-11*, *cox-15*, *cox-17*, *cyc-1*, *F45H10.2*, *F58F12.1*, *gas-1*, *hpo-18*, *isp-1*, *lpd-5*, *nduf-5*, *nduf-6*, *nduf-7*, *nuo-1*, *nuo-2*, *nuo-4*, *nuo-6*, *R04F11.2*, *R07E4.3*, *R53.4*, *T02H6.11*, *T27E9.2*, *ucr-2.2*, *ucr-11*, *vha-1*, *vha-2*, *vha-4*, *vha-6*, *vha-8*, *vha-9*, *vha-10*, *vha-12*, *vha-14*, *vha-16*, and *Y69A2AR.18* (Data S17). Remarkably, 48 genes involved in the oxidative phosphorylation pathway were significantly down-regulated due to mutation in *acdh-12*, which might be indicative of its function as an assembly factor for mitochondrial complex I ^46^. Oxidative phosphorylation impairment induced by the 48 genes suppression could be detrimental for mitochondrial respiration and energy supply, which might contribute to the reduction of salinity-stress tolerance induced by *acdh-12* mutation.

We found that the significantly up-regulated genes in the autophagy pathway in *osm-11;acdh-12* double mutants included *alfa-1*, *allo-1*, *atg-2*, *atg-9*, *atg-13*, *atg-16.1*, *atg-18*, *C35E7.2*, *C35E7.4*, *ceh-86*, *dct-1*, *epg-4*, *epg-5*, *F44F1.4*, *hpk-1*, *ikke-1*, *ZK1053.3*, and *ZK1053.4* (Data S20). However, in many cases, increased autophagy promotes cell survival, but unlimited autophagy induces oxidative stress-related cell death ^57^. Autophagy of mitochondrion, known as mitophagy, is a specialized form of autophagy that maintains cellular homeostasis by eliminating damaged mitochondria ^58,59^. Recent studies have shown that excessive mitophagy can lead to cell death through both apoptosis and ferroptosis ^60^. Under 30‰ salinity, compared to *osm-11* single mutants, the significantly up-regulated genes in the “autophagy of mitochondrion” GO term in *osm-11;acdh-12* double mutants included *atg-2*, *atg-9*, *atg-13*, *atg-18*, and *dct-1* (Data S20). *dct-1* encodes a mitophagy receptor, we found that RNAi knockdown *of dct-1* significantly increased the survival rate of *osm-11;acdh-12* double mutants, under 30‰ salinity condition, suggesting that *acdh-12* mutation significantly reduced salinity-stress tolerance of *osm-11* mutants through mitophagy.

Interestingly, under 30‰ salinity, compared to *osm-11* single mutants, the significantly down-regulated genes in glutathione metabolism pathway in *osm-11;acdh-12* double mutants contained *gstk-1*, *T03D8.6*, *gst-41*, *txdc-12.1*, *rnr-2*, *idh-1*, *gst-1*, *gst-4*, *gst-5*, *gst-6*, *gst-7*, *gst-10*, *gst-38*, *gspd-1*, *gst-36*, *spds-1*, *gsto-1*, *E01A2.1*, *gpx-1*, *gpx-3*, *gpx-5*, *gpx-7*, *C44B7.7*, *T25B9.9*, and *gst-25* (Figure 4C; Data S17). Of these genes, both *gpx-1* and *gpx-7* encode orthologs of human GPX4 (glutathione peroxidase 4), which is a key selenoenzyme protecting against ferroptosis by converting toxic lipid hydroperoxides into non-toxic lipid alcohols ^61^. Additionally, we found that the expressions of *ftn-1*, *ads-1*, *dhod-1*, as well as *mboa-6* and *mboa-7* was significantly repressed in *osm-11;acdh-12* double mutants compared to *osm-11* single mutants, under salinity-stress conditions. Intriguingly, we found that RNAi knockdown of *ftn-1* did not lead to a significant reduction of survival rate in *osm-11* mutants. However, we observed that RNAi knockdown of *gpx-1* and *gpx-7* markedly decreased the survival of *osm-11* mutants under hyper-saline conditions, indicating that salinity-stress tolerance in *osm-11* mutants might be due to higher GPX4 homolog genes’ expression and their ability to convert toxic lipid hydroperoxides into non-toxic lipid alcohols, which might lead to ferroptosis.

LDs are cellular organelles that dynamically regulate lipid and energy homeostasis ^47^. We found that compared to *osm-11* mutants, the expression of short-chain dehydrogenase/reductase encoding gene *dhs-28* and sterol carrier protein encoding gene *daf-22* was significantly down-regulated in *osm-11;acdh-12* double mutants. Moreover, we showed that the hyper-saline resistance of *osm-11;daf-22* and *osm-11;dhs-28* mutants was significantly reduced compared to *osm-11* single mutants, suggesting that salinity-stress tolerance of *osm-11* mutants may depend on *dhs-28* and *daf-22*, in line with mutations in *dhs-28* and *daf-22* led to LD enlargement ^47^.

Our findings highlight the critical roles of *osm-11* and *acdh-12* in modulating *C. elegans* to tolerate salinity stress. *acdh-12* mutation disrupted mitochondrial homeostasis in *osm-11* mutants, significantly impairing their salinity tolerance and underscoring the essentiality of mitochondrial stability for survival under hyper-saline conditions. The *osm-11* mutants likely enhanced salt tolerance by mitigating mitochondrial fragmentation, mitophagy, and ferroptosis partially through ACDH-12. Additionally, salinity tolerance in *osm-11* mutants relies on *gpdh-1/2* (involved in glycerol biosynthesis) and *dhs-28/daf-22* (key genes in mitochondrial lipid β-oxidation), indicating coordinated metabolic pathways. Mechanistically, we found that *osm-11* mutants promote hyper-saline tolerance parallel with insulin-like signaling in a canonical Notch signaling pathway (see Section 9). Our study opens new avenues for investigating how soil organisms adapt to increased salinity levels, providing insights into broader potentials of adaptive mechanisms in the context of global climate change. These findings provide novel insights into soil organisms’ adaptive strategies to salinity stress, offering theoretical foundations for understanding ecosystem resilience and guiding conservation efforts in salt-impacted terrestrial habitats.

## Acknowledgements

We thank all the members of the L.Z lab for their valuable insights and discussions. Gratitude also goes out to Profs. Jianhai Xiang, Guofan Zhang, Fuhua Li, Li Li, Li Sun, Guangce Wang, Chaomin Sun, and Baozhong Liu’s help for suggestions, reagents and facilities.

## Funding

This work was supported by the National Natural Science Foundation of China [No. 32170537]; Key deployment project of Centre for Ocean Mega-Research of Science, Chinese Academy of Sciences [COMS2019Q16]; “Talents from overseas Program, IOCAS” of the Chinese Academy of Sciences; and “Qingdao Innovation Leadership Program” [Grant 16-8-3-19-zhc]. Some nematode strains were provided by the Caenorhabditis Genetics Center, which is funded by NIH Office of Research Infrastructure Programs [P40 OD010440].

## Notes

### Competing Interest Statement

The authors have declared no competing interest.

